# Nrf2 controls homeostatic transcriptional signatures and inflammatory responses in a cell-type specific manner in the adult mouse brain

**DOI:** 10.1101/2024.11.14.623657

**Authors:** Xin He, Owen Dando, Jing Qiu

**Author notes:** Correspondence to Jing Qiu.

## Abstract

Nrf2 is an attractive potential therapeutic target for various neurological disorders including neurodegenerative diseases. The mechanisms behind Nrf2-mediated cytoprotection are incompletely understood, however, an anti-inflammatory effect is a popular hypothesis, based primarily on studies where the prior pharmacological activation of Nrf2 has suppressed subsequent responses to inflammatory insults. In the adult brain, Nrf2 is highly expressed in microglia, astrocytes and brain endothelial cells, with minimal expression in neurons. As yet, the brain cell type-specific role of Nrf2 in regulating the basal transcriptome and controlling neuroinflammation is unknown. To address this, we employed three inducible conditional Nrf2 knockout mice in which Nrf2 is deleted in microglia, astrocytes, and brain endothelial cells respectively in a tamoxifen-inducible manner. We discovered that in the healthy brain Nrf2 controls distinct transcriptional signatures in the three brain cell types under study. Surprisingly, following a systemic inflammatory challenge, we found that Nrf2-deficient microglia and astrocytes (but not endothelial cells) both mount a suppressed inflammatory response compared to wild type cells. Moreover, comparison of Nrf2-deficient with wild-type microglia revealed that even in the absence of any insult, microglial Nrf2 supports basal activity of inflammation response genes. Thus, our study uncovers a novel cell type-specific role for endogenous Nrf2 activity in supporting inflammatory responses in the brain.

## Introduction

The transcription factor Nrf2 (encoded by the *Nfe2l2* gene) is a widely-expressed stress-responsive master regulator of genes involved in important aspects of homeostatic physiology^1,2,3^. Under normal conditions Nrf2 is targeted for ubiquitin-mediated degradation by Keap1, but in response to stress, this interaction is inhibited, leading to Nrf2 accumulation in the nucleus where it regulates the expression of genes that contain antioxidant response elements (AREs) in their promoter^1,2^. In the brain, Nrf2 is highly expressed in microglia, astrocytes and brain endothelial cells (BECs), with low expression in neurons^4^. However, the cell-type specific roles of Nrf2 in the brain was not well-understood, partly due to a historical reliance on global Nrf2 knockout mice to probe the roles of Nrf2^5^. This knowledge gap is important to fill because cell type-specific roles of Nrf2 are likely to be influenced by the function of that cell type, and from a translational point of view one ideally needs to know the locus of action of any Nrf2-targeting therapeutic to optimise bioavailability.

Previous studies have highlighted an anti-inflammatory role of Nrf2 in the brain, via possible mechanisms such as elimination of reactive oxygen species (ROS), blocking proinflammatory cytokine transcription, or reducing expression of adhesion molecules^6,7,8,9^. However, most of these studies were based on prior pharmacological activation of Nrf2 using natural or synthetic compounds of Nrf2 activators^9^. The role of the endogenous activity of Nrf2 in controlling inflammatory responses in individual brain cell types is not clear. The global Nrf2 knockout mouse has pleiotropic phenotypes in development and maturity^10^, which, in addition to the non-cell autonomous effects between different cell types, makes it difficult to dissect the cell-type specific role of Nrf2 in the brain under inflammatory conditions. In this study, we hypothesized that Nrf2 plays distinct cell type-specific roles both in controlling the basal transcriptome as well as determining those cell types’ responses to inflammatory insults.

## Results

### Generation of inducible-conditional Nrf2 knockout mice in microglia and astrocytes

To investigate the cell-type specific role of Nrf2 in the brain, we used *Nefe2l2*^*fl/fl*^ mice which possess *loxP* sites flanking exon 5 of the *Nefe2l2* gene which encodes the DNA binding domain of Nrf2. Previously, we generated an inducible-conditional EC-specific knockout line (*Nfe2l2*^ENDO^) mouse by crossing *Nefe2l2*^*fl/fl*^ mice with *Cdh5*^*CreERT*28^ that expresses tamoxifen-inducible CreERT specifically in endothelial cells^11^. In the current study, we crossed *Nefe2l2*^*fl/fl*^ mice with either an inducible astrocyte-specific Cre mouse (*ALDH1L1*^*CreERT*^) or an inducible monocyte-specific Cre mouse line (*CX3CR1*^*CreERT*^). Following tamoxifen injection, exon 5 of the Nrf2 gene is deleted exclusively either in astrocytes to generate an inducible-conditional astrocyte-specific knockout line (*ALDH1L1*^*CreERT2*^:*Nfe2l2*^*fl/fl*^, hereafter *Nfe2l2*^Astr^), or exclusively in microglia to generate an inducible-conditional monocyte-specific knockout line (*CX3CR1*^*CreERT2*^:*Nfe2l2*^*fl/fl*^, hereafter *Nfe2l2*^Micr^). Tamoxifen injection will cause deletion of exon 5 of Nrf2 in both microglia and blood monocyte-derived macrophages, but within 4 weeks blood monocytes turn over^11^, leaving only microglia (half-life approximately 2 years) lacking exon 5 of Nrf2. Sorting of brain cells by FACS followed by qPCR confirmed the successful deletion of Nrf2 exon 5 in either microglia in *Nfe2l2*^Micr^ mice or astrocytes in *Nfe2l2*^Astro^ mice without affecting Nrf2 exon 5 levels in other brain cell types in either mouse line (Fig. 1A & B).

**Fig. 1.**
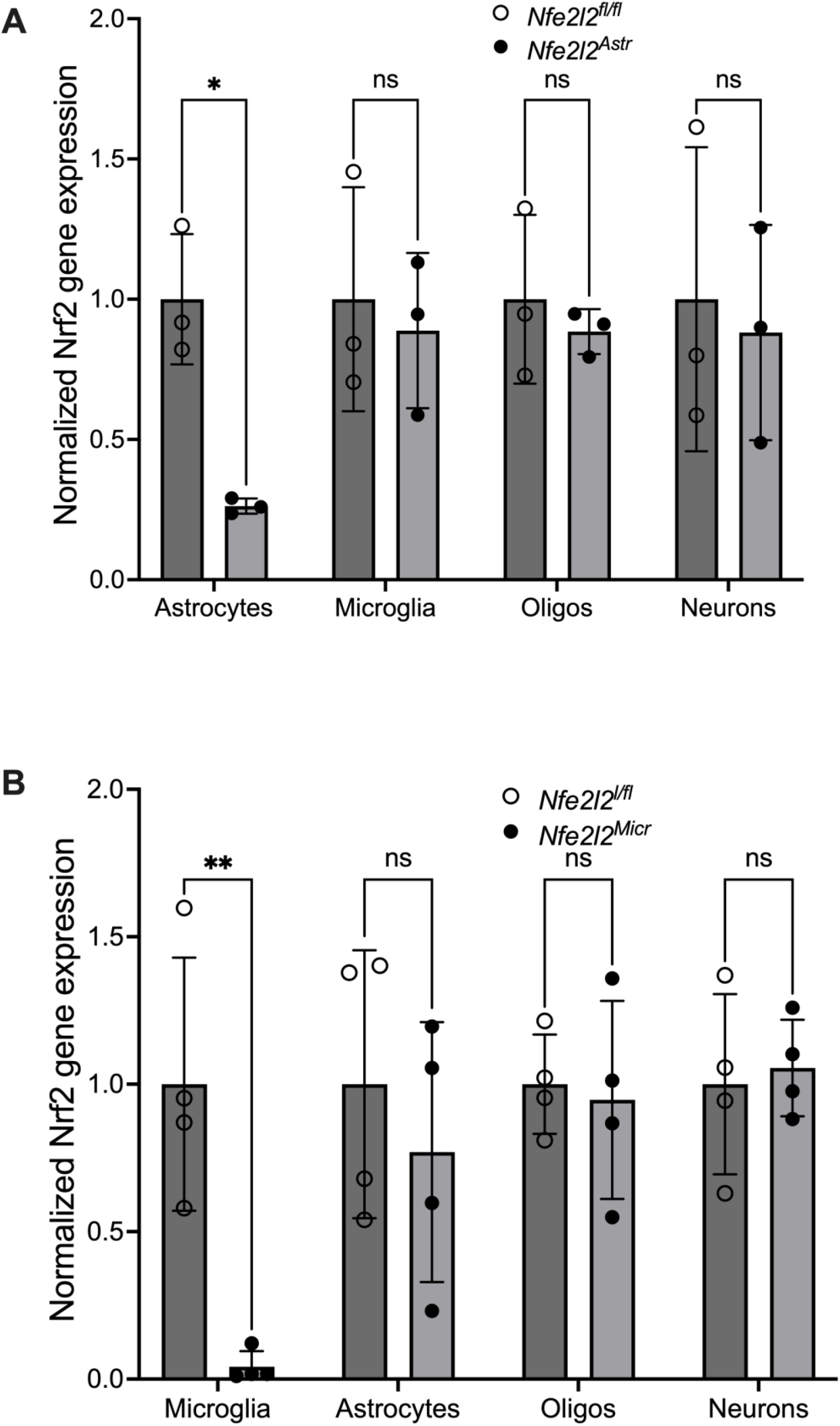
Generation of inducible-conditional Nrf2 knockout mice in microglia and astrocytes. **(A)** qRT-PCR confirms gene expression of exon 5 of Nrf2 is specifically deleted in astrocytes, but not in microglia, oligodendrocytes, or neurons in *Nfe2l2*^ASTR^ mice. From left to right p=0.049, 0.9890, 0.9875, and 0.9862, n=4, unpaired t-test. **(B)** qRT-PCR confirms gene expression of exon 5 of Nrf2 is specifically deleted in microglia, but not in astrocytes, oligodendrocytes, or neurons in *Nfe2l2*^MICR^ mice. From left to right p=0.0014, 0.7955, 0.9989, and 0.9988, n=4, unpaired t-test.

### Nrf2 controls distinct transcriptional signatures in microglia, astrocytes and BECs

Previously, using the *Cdh5*^*CreERT*2^ mice, we found that Nrf2 is required to maintain a BEC transcriptional signature in the adult mouse brain^11^. In this current study, we have extended this analysis to microglia from *Nfe2l2*^Micr^ mice and astrocytes from *Nfe2l2*^Astr^ mice. RNA-seq analysis of differentially regulated genes in Nrf2-deficient microglia from *Nfe2l2*^Micr^ mice revealed 33 upregulated and 24 downregulated genes (p<0.05; Fig. 2A). NRF2-deficient microglia exhibited a down-regulation of known Nrf2 target genes (*Hmox1 and Srxn1*, Fig. 2A). Functional categorisation using the Enrichr Bioinformatic database for GO term enrichment analysis of biological process (BP), molecular function (MF), and KEGG pathway enrichment indicate that Nrf2 controls key transcriptional signatures in microglia (Table 1). For example, Nrf2-deficient microglia have dysregulated immunity and iron ion homeostasis, reduced response to oxidative stress, chemical stress, cytokine stimulus and bacterium, and dysregulated protein folding. Cellular compartment (CC) enrichment analysis revealed reduced expression of secretory granule and phagocytic vesicle membrane. KEGG pathway suggested downregulation of genes involved in infection and neurodegenerative diseases.

**Table 1.**
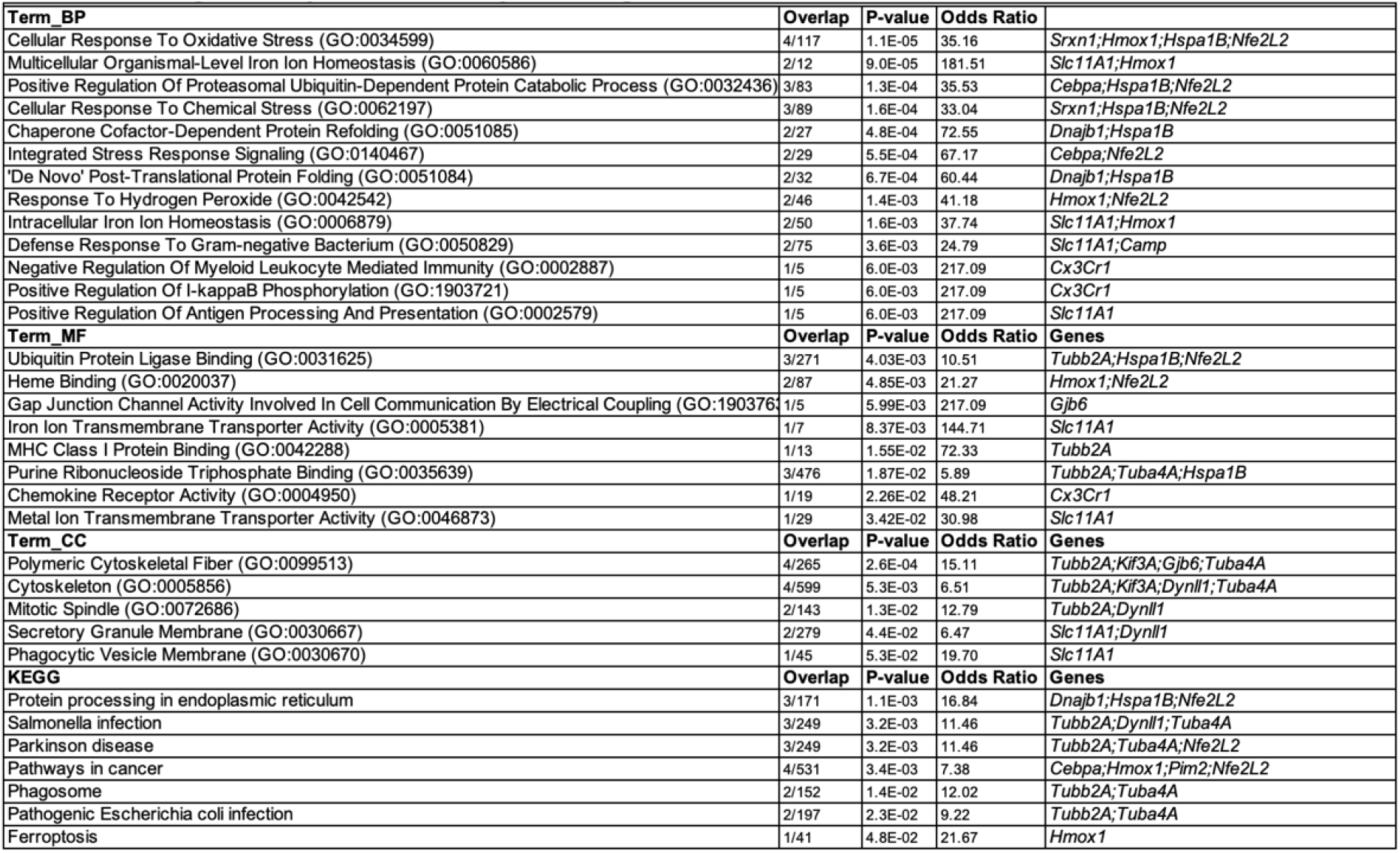
Genes Downregulated by Nrf2-deficiency in microglia in *Nfe212*^*Micr*^ mice under basal condition.

**Fig. 2.**
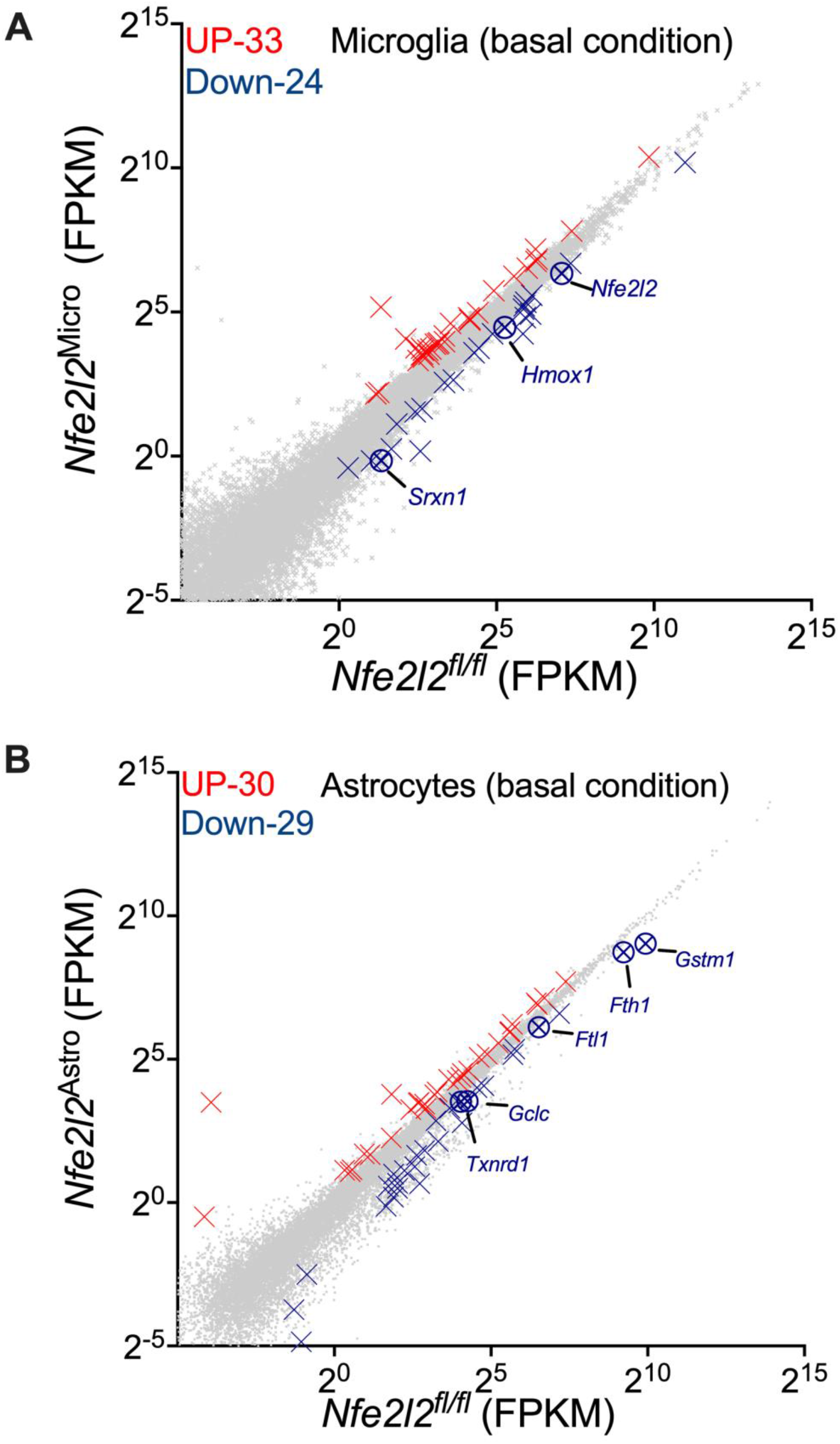
Nrf2 maintains homeostatic transcriptional signatures in microglia and astrocytes. Scatterplot of RNA-seq analysis was generated for genes with average expression >0.1 FPKM across the data sets. Highlighted with red and blue crosses are the genes whose expression are significantly increased or decreased respectively (*DESeq2 P*_adj<0.05, n=4-6). **(A)** The influence of Nrf2 specific KO in microglia on microglial transcriptome in *Nfe2l2*^MICR^ mice at basal condition. **(B)** The influence of Nrf2 specific KO in astrocytes on astrocyte transcriptome in *Nfe2l2*^ASTR^ mice at basal condition.

Analysis of differentially regulated genes in Nrf2-deficient astrocytes in in *Nfe2l2*^Astr^ mice revealed 30 upregulated and 29 downregulated genes (p<0.05; Fig. 2B). As expected, Nrf2-deficient astrocytes in *Nfe2l2*^Astr^ mice exhibited a down-regulation of several known Nrf2 target genes (e.g. *Gstm1, Fth1, Ttl1, Gclc*, and *Txnrd1*; Fig. 2B). Functional categorisation using GO term enrichment analysis of BP, MF, and KEGG pathways suggest that Nrf2-deficient astrocytes have reduced expression of genes involved in glutathione metabolism, vitamin B6 metabolism, nicotinate and nicotinamide metabolism, and tyrosine metabolism. Nrf2-deficient astrocytes exhibited a reduced expression of genes involved in the response to oxidative stress, ferroptosis, and axon regeneration (Table 2). CC enrichment analysis revealed decreased expression of genes in lysosome, peroxisome, and perisynatic extracellular matrix regions (Table 2).

**Table 2.**
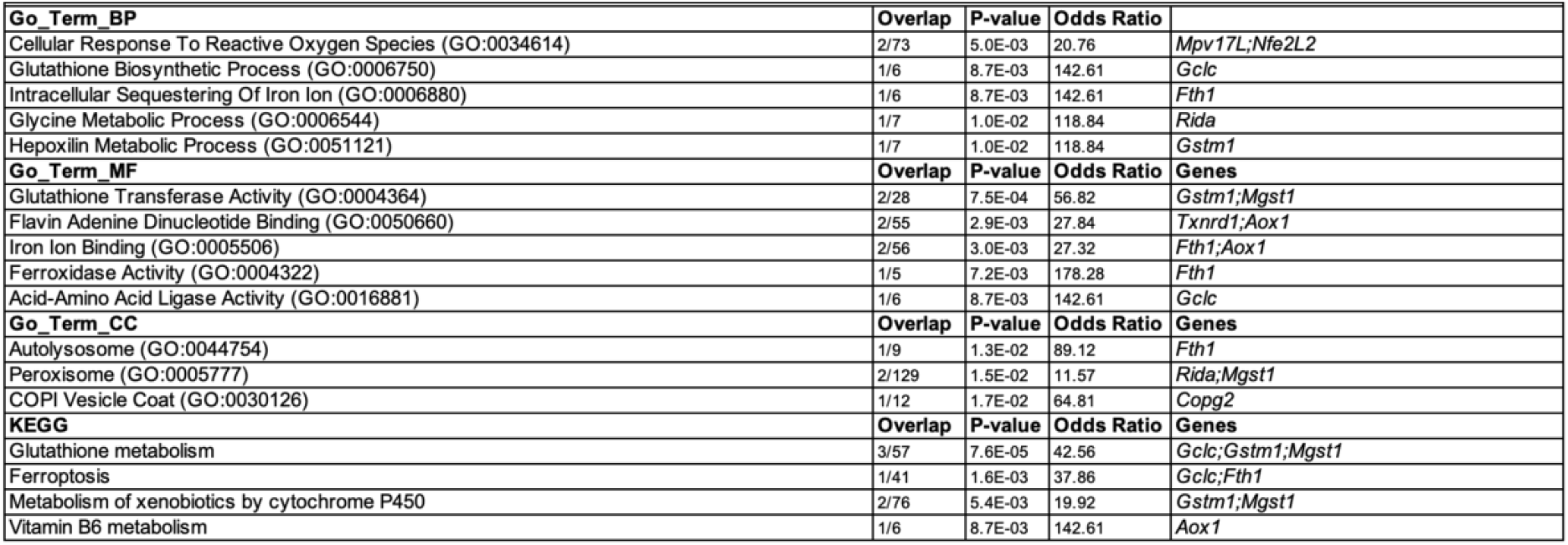
Genes Downregulated by Nrf2-deficiency in astrocytes in *Nfe212* ^*Astr*^ mice under basal condition.

Previously, using an inducible conditional-endothelial Nrf2 KO mice (*Nfe2l2*^Endo^), we found that basal Nrf2 activity controls key transcriptional signatures in BECs as well^11^. In this study, we compared the differentially regulated genes under basal conditions in microglia in Nfe2l2^Micr^ with those in astrocytes in *Nfe2l2*^Astr^, and in BECs in *Nfe2l2*^Endo^ (Fig. 3A & B), we found only a few genes common between astrocytes and BECs. These data suggest that Nrf2 controls distinct transcriptional programmes in these three cell types under basal conditions.

**Fig. 3.**
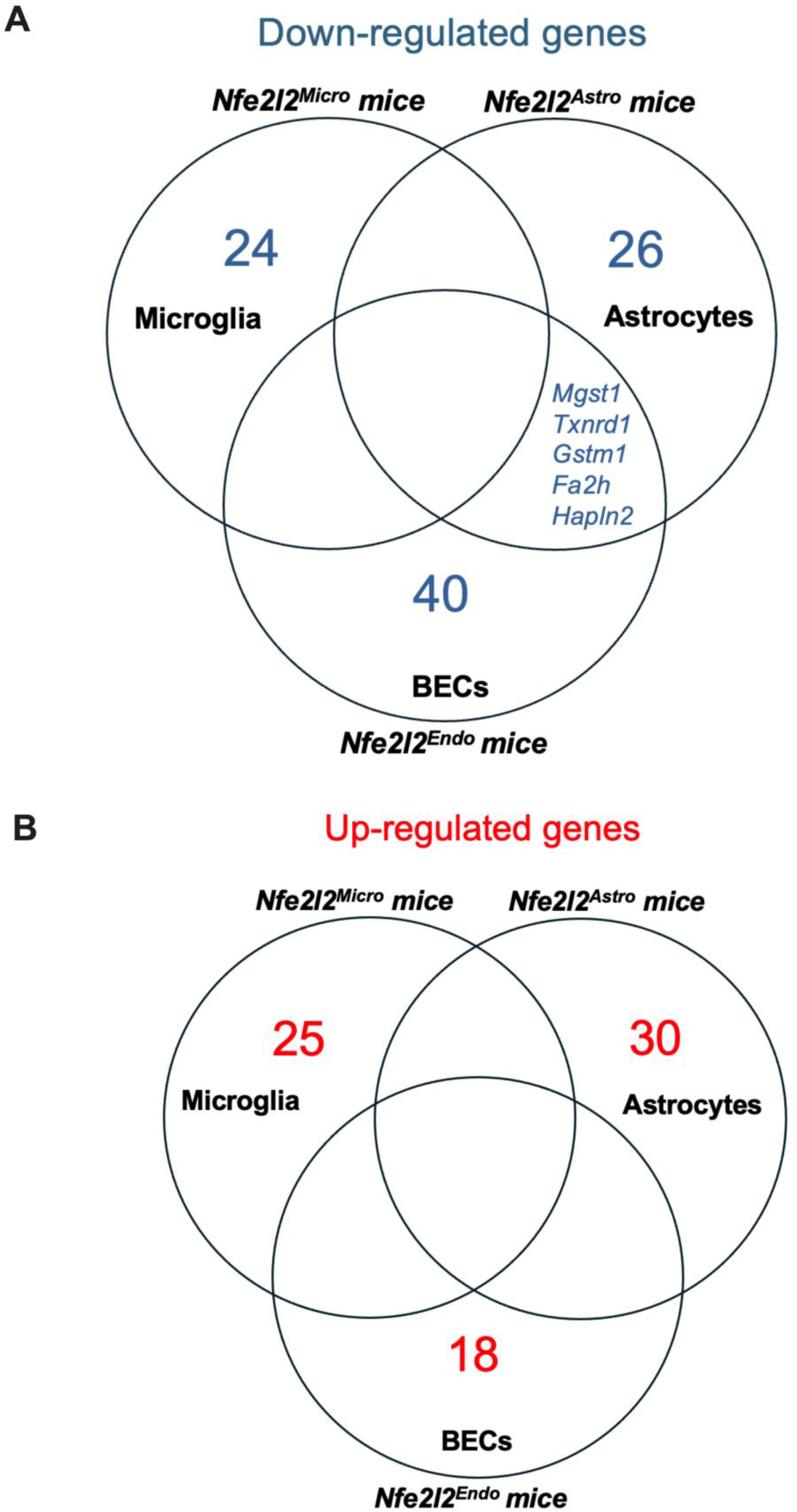
Nrf2 controls distinct transcriptional signatures in microglia, astrocytes and BECs. **(A)** The significantly downregulated genes due to Nrf2 KO in microglia in *Nfe2l2*^MICR^ mice, in astrocytes in *Nfe2l2*^ASTR^ mice, and in BECs in *Nfe2l2*^ASTR^ mice, compared to the respective littermate controls, were compared using Venn diagram. **(B)** The significantly upregulated genes due to Nrf2 KO in microglia in *Nfe2l2*^MICR^ mice, in astrocytes in *Nfe2l2*^ASTR^ mice, and in BECs in *Nfe2l2*^ASTR^ mice, compared to the respective littermate controls, were compared using Venn diagram.

### Nrf2 controls inflammatory responses in microglia, astrocytes and BECs following systemic inflammation

We next wanted to determine the impact of Nrf2 deficiency on cellular inflammatory responses. It is well-established that systemic inflammation can lead to blood brain barrier (BBB) compromise and neuroinflammation, so we employed a model of systemic inflammation to determine how Nrf2 status affects different brain cells’ response to inflammatory insults. We induced systemic inflammation in mice by intraperitoneal injection of bacterial endotoxin LPS in *Nfe2l2*^LoxP/LoxP^ mice, chosen as a “conditional-ready” control to compare with conditional knockout mice used in the study. We showed in our previous study that microglia, astrocytes and BECs display widespread transcriptional changes following LPS exposure in *Nfe2l2*^LoxP/LoxP^ mice^11^. Here we wanted to know how Nrf2 deficiency influenced the transcriptome of microglia, astrocytes and BECs under these po-inflammatory conditions. RNAseq analysis of Nrf2-deficient microglia in *Nfe2l2*^Micr^ mice revealed 56 upregulated and 116 downregulated genes compared to *Nfe2l2*^LoxP/LoxP^ microglia in mice treated with LPS insults (p<0.05; Fig. 4A). Functional categorisation using KEGG pathway and GO term enrichment analysis of BP (Table 3) suggests that pathways involved in leukocyte transendothelial migration, response to cytokine and response to Tumour Necrosis Factor were reduced in *Nfe2l2*^Micr^ microglia. Analysis of Nrf2-deficient astrocytes in *Nfe2l2*^Astr^ mice revealed 103 upregulated and 128 downregulated genes compared to *Nfe2l2*^LoxP/LoxP^ astrocytes in mice treated with LPS insults (p<0.05; Fig. 4B). Functional categorisation using KEGG pathway and GO term enrichment analysis of BP (Table 4) suggests that inflammatory pathways including MAPK signalling pathway, JAK-STAT signalling pathway, and p53 signalling pathway were downregulated due to Nrf2-deficiency in astrocytes following systemic LPS insults. In addition, Wnt signalling pathway, HIF-1 signalling pathway, and ubiquitin mediated proteolysis were also downregulated, suggesting Nrf2-deficiency astrocytes reduced responses to inflammatory stimulus. In contrast to Nrf2-deficient microglia and astrocytes, Nrf2-deficient BECs in *Nfe2l2*^Endo^ mice had modest number of differentially regulated genes in response to peripheral LPS compared to *Nfe2l2*^LoxP/LoxP^ mice (Fig. 4C). Only 66 genes were differentially expressed in *Nfe2l2*^Endo^ BECs compared to *Nfe2l2*^LoxP/LoxP^ astrocytes following LPS treatment. Thus, endogenous Nrf2 activity has a stronger influence on inflammatory responses in microglia and astrocytes than in BECs.

**Table 3.** Genes Downreagulated by Nrf2 – deficiency in microglia in *Nfe212*^*MIcr*^ mice under inflammatory condition.

| KEGG | Overlap | P-value | Odds Ratio | Genes |
| --- | --- | --- | --- | --- |
| Cell adhesion molecules | 6/148 | 2.3E-04 | 7.58 | Cdh5;Cldn5;Esam;Nrcam;Ncam2;Sele |
| Fatty acid elongation | 2/27 | 1.1E-02 | 13.94 | Echs1;Acot1 |
| Butanoate metabolism | 2/28 | 1.1E-02 | 13.40 | Echs1;Abat |
| beta-Alanine metabolism | 2/30 | 1.3E-02 | 12.44 | Echs1;Abat |
| ECM-receptor interaction | 3/88 | 1.5E-02 | 6.18 | Col4A2;Col4A1;Spp1 |
| Leukocyte transendothelial migration | 3/114 | 2.9E-02 | 4.73 | Cdh5;Cldn5;Esam |
| Pyruvate metabolism | 2/47 | 3.0E-02 | 7.73 | Pkm;Akr1A1 |
| Term_BP | Overlap | P-value | Odds Ratio | Genes |
| Response To Cytokine (GO:0034097) | 8/125 | 6.7E-07 | 12.51 | Csf3;Sphk1;Mx1;Timp3;Ccr7;Timp1;Sele |
| Response To Tumor Necrosis Factor (GO:0034612) | 5/108 | 4.2E-04 | 8.65 | Dhx9;Sphk1;Gbp2;Sele;Gbp3 |
| Response To Lipid (GO:0033993) | 5/110 | 4.5E-04 | 8.49 | Acer2;Spp1;Sox9;Trim16;Sele |
| Positive Regulation Of Phosphatidylinositol 3-Kinase Signaling (GO:0014068) | 4/89 | 6.9E-04 | 10.89 | Csf3;Nedd4;Sox9;Dcn |

**Table 4.**
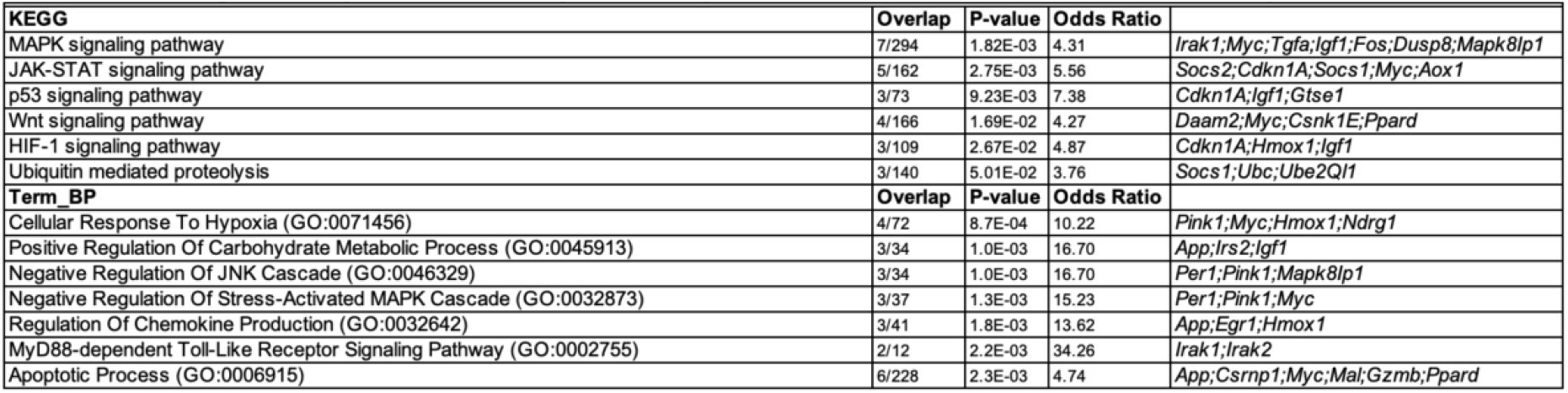
Genes Downreagulated by Nrf2 – deficiency in astrocytes in *Nfe212*^*Astr*^ mice under inflammatory condition.

**Fig. 4.**
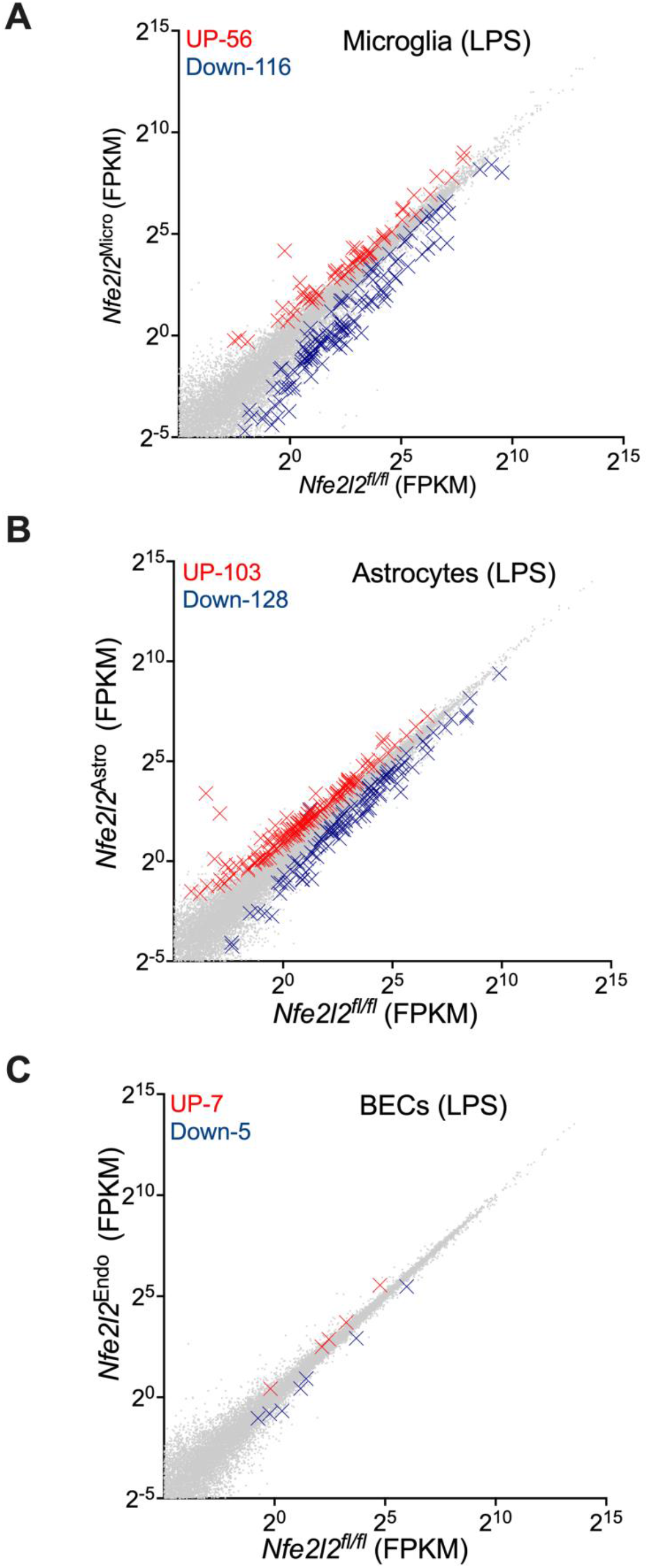
Nrf2 controls inflammatory responses in microglia, astrocytes and BECs under systemic inflammation. Mice were given i.p. injection of either saline or LPS for 24 hrs. The brain cells were isolated and then sorted with FACS for microglia, astrocytes and BECs, RNA extracted and RNA-seq performed. Scatterplot was generated for genes with average expression >0.1 FPKM across the data sets. Highlighted with red and blue crosses are the genes whose expression are significantly increased or decreased respectively (*DESeq2 P*_adj<0.05, n=6-8). ‘N’ refers here and throughout as an independent replicate (i.e. a mouse). **(A)** Scatterplot of RNA-seq analysis showing the microglial transcriptome modified by Nrf2 KO in *Nfe2l2*^MICR^ mice compared to *Nfe2l2*^*fl/fl*^ mice in response to peripheral LPS insults. **(B)** Scatterplot of RNA-seq analysis showing the astrocyte transcriptome modified by Nrf2 KO in astrocytes in *Nfe2l2*^ASTR^ mice compared to *Nfe2l2*^*fl/fl*^ mice in response to peripheral LPS insults. **(C)** Scatterplot of RNA-seq analysis showing the BEC transcriptome modified by Nrf2 KO in BECs in *Nfe2l2*^ENDO^ mice compared to *Nfe2l2*^*fl/fl*^ mice in response to peripheral LPS insults.

### After a systemic insult, Nrf2 supports the proinflammatory responses in microglia and astrocytes, but not in BECs

We next wanted to determine whether the groups of genes up or down regulated following LPS treatment show any pattern of perturbation due to Nrf2 deficiency. We firstly assessed whether Nrf2 was able to modulate microglial overall responses to LPS in *Nfe2l2*^Micr^ mice. We took microglial genes up- and down-regulated by LPS and studied the impact of Nrf2-deficiency in microglia. We found that the gene set induced by LPS in microglia was overall repressed as the result of Nrf2-deficiency in microglia (Fig. 5A & G), and the gene set repressed by LPS in microglia was overall elevated as the result of Nrf2-deficiency in microglia (Fig. 5B & H). This suggests that Nrf2-deficiency in microglia dampened down microglial proinflammatory responses to peripheral LPS insults. Using the same strategy, we assessed the influence of basal Nrf2 activity on modulating astrocytes and BEC responses to LPS in *Nfe2l2*^Astr^ mice and *Nfe2l2*^Endo^ mice respectively. We found that similar to Nrf2-deficient microglia, the gene set induced by LPS in astrocytes was overall repressed as the result of Nrf2-deficiency microglia (Fig. 5B & G), and the gene set repressed by LPS in astrocytes was overall elevated as the result of Nrf2-deficiency in astrocytes (Fig. 5C & H). This suggests that like microglia, Nrf2-deficiency in astrocytes reduced astrocytes’ overall proinflammatory responses to peripheral LPS insults. In contrast, the gene sets induced or repressed by LPS in BECs were not overall changed as the result of Nrf2-deficiency in BECs in *Nfe2l2*^Endo^ mice (Fig. 5E & G; F & H). Thus, basal Nrf2 activity in microglia and astrocytes overall enhances the responses of these cell type to peripheral inflammation, contrary to the classical view of Nrf2 being solely anti-inflammatory.

**Fig. 5.**
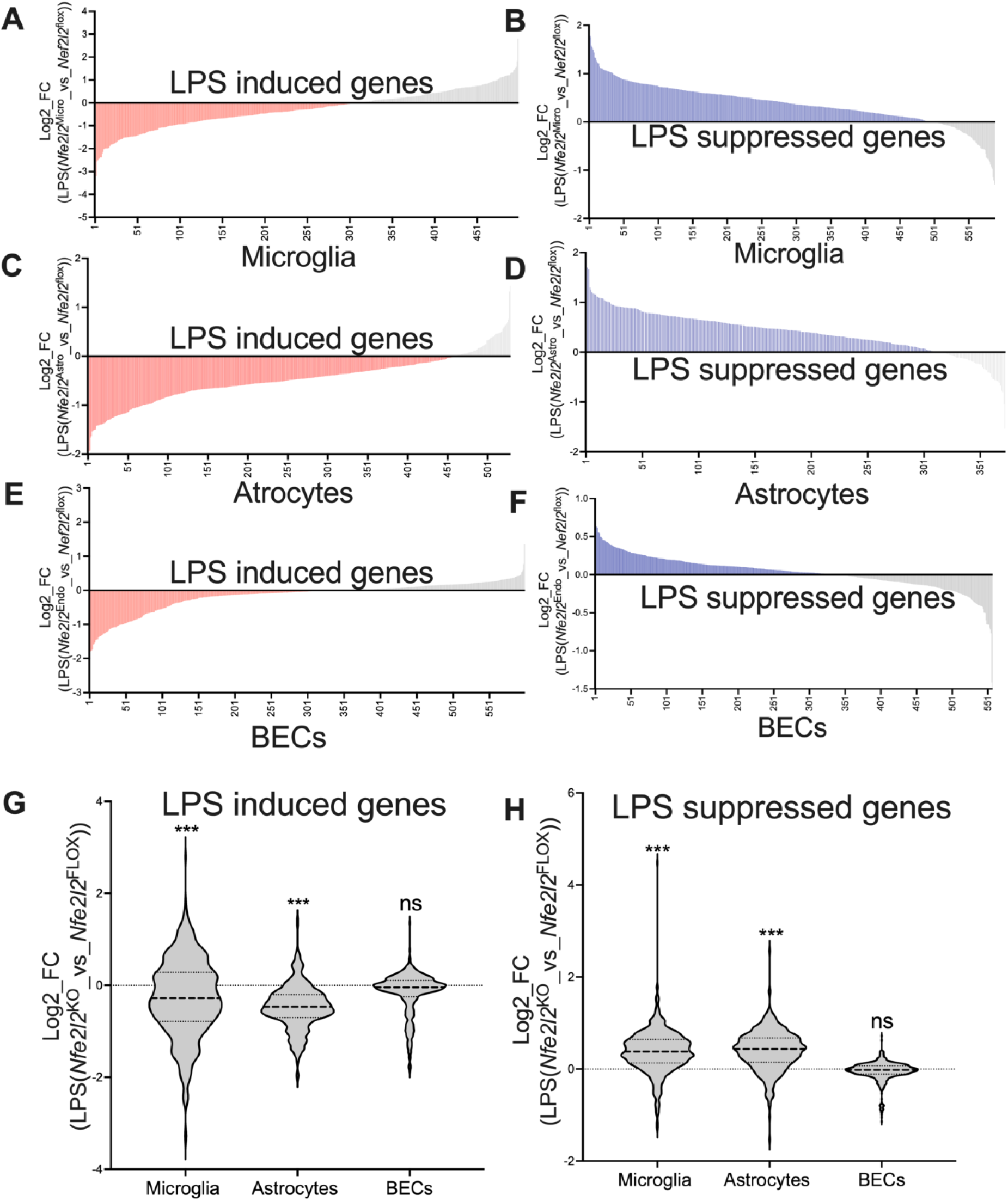
Nrf2 promotes proinflammatory responses in microglia and astrocytes, but not in BECs under systemic inflammation. **(A)** The influence of Nrf2 KO in microglia on the expression of microglial LPS-induced gene set under inflammatory condition. For microglial LPS-induced genes (x-axis, FPKM>1, *DESeqP*_adj<0.05, Log_2_FC >1) in WT mice, the Log_2_FC (y-axis) of each gene (x-axis) due to Nrf2 KO in microglia in response to peripheral LPS insults were plotted. **(B)** The influence of Nrf2 KO in microglia on the expression of microglial LPS-repressed gene set under inflammatory condition. For microglial LPS-repressed genes (x-axis, FPKM>1, *DESeqP*_adj<0.05, Log_2_FC >1) in WT mice, the Log_2_FC (y-axis) of each gene (x-axis) due to Nrf2 KO in microglia in response to peripheral LPS insults were plotted. **(C)** The influence of Nrf2 KO in astrocytes on the expression of astrocyte LPS-induced gene set under inflammatory condition. For astrocyte LPS-induced genes (x-axis, FPKM>1, *DESeqP*_adj<0.05, Log_2_FC >1) in WT mice, the Log_2_FC (y-axis) of each gene (x-axis) due to Nrf2 KO in astrocytes in response to peripheral LPS insults were plotted. **(D)** The influence of Nrf2 KO in astrocytes on the expression of astrocyte LPS-repressed gene set under inflammatory condition. For astrocyte LPS-repressed genes (x-axis, FPKM>1, *DESeqP*_adj<0.05, Log_2_FC >1) in WT mice, the Log_2_FC (y-axis) of each gene (x-axis) due to Nrf2 KO in astrocytes in response to peripheral LPS insults were plotted. **(E)** The influence of Nrf2 KO in BECs on the expression of BEC LPS-induced gene set under inflammatory condition. For BEC LPS-induced genes (x-axis, FPKM>1, *DESeqP*_adj<0.05, Log_2_FC >1) in WT mice, the Log_2_FC (y-axis) of each gene (x-axis) due to Nrf2 KO in BECs in response to peripheral LPS insults were plotted. **(F)** The influence of Nrf2 KO in BECs on the expression of BEC LPS-repressed gene set under inflammatory condition. For BEC LPS-repressed genes (x-axis, FPKM>1, *DESeqP*_adj<0.05, Log_2_FC >1) in WT mice, the Log_2_FC (y-axis) of each gene (x-axis) due to Nrf2 KO in BECs in response to peripheral LPS insults were plotted. **(G)** From left to right, *p=4.9E-12, F (1,3859) =47.81 relates to main effect of Nrf2-KO on the expression of microglial LPS-induced gene set under inflammatory conditions, 2-way ANOVA; *p=1.5E-05, F (1,4761) =18,76 relates to main effect of Nrf2-KO on the expression of astrocyte LPS-induced gene set under inflammatory condition, 2-way ANOVA; p=0.5382, F (1,3594) =0.3789 relates to main effect of Nrf2-KO on the expression of BEC LPS-induced gene set under inflammatory condition, 2-way ANOVA **(H)** From left to right, *p=1.1E-31, F (1,6993) =138.6 relates to main effect of Nrf2-KO on the expression of microglial LPS-repressed gene set under inflammatory condition, 2-way ANOVA; *p=6.1E-18, F (1,3357) =75.34 relates to main effect of Nrf2-KO on the expression of astrocyte LPS-repressed gene set under inflammatory condition, 2-way ANOVA; p=0.5382 F (1,5570) =0.9470 relates to main effect of Nrf2-KO on the expression of BEC LPS-repressed gene set under inflammatory condition, 2-way ANOVA

### Nrf2 mediates proinflammatory gene expression in microglia, not in astrocytes and BECs at basal condition

Finally, we wanted to know whether the impact of Nrf2-deficiency controls the previously described LPS response genes under basal, non-inflammatory conditions. We found that the gene set induced by LPS in microglia was overall repressed as the result of Nrf2-deficiency in microglia under basal conditions (Fig. 6A & G), and the gene set repressed by LPS in microglia was overall elevated as the result of Nrf2-deficiency in microglia under basal conditions (Fig. 6B & H). In contrast, when we took the gene sets of either up- or down-regulated by peripheral LPS insults in astrocytes or BECs and studied the impact of Nrf2-deficiency on expression of these genes in these two cell types under basal conditions, we found that Nrf2-deficiency had no impact on the expression of these gene sets in either astrocytes (Fig. 6C & G; D & H) or BECs (Fig. 6E & G; F & H). Overall these data suggest that Nrf2 controls the basal expression of inflammatory response genes in a cell-type specific manner.

**Fig. 6.**
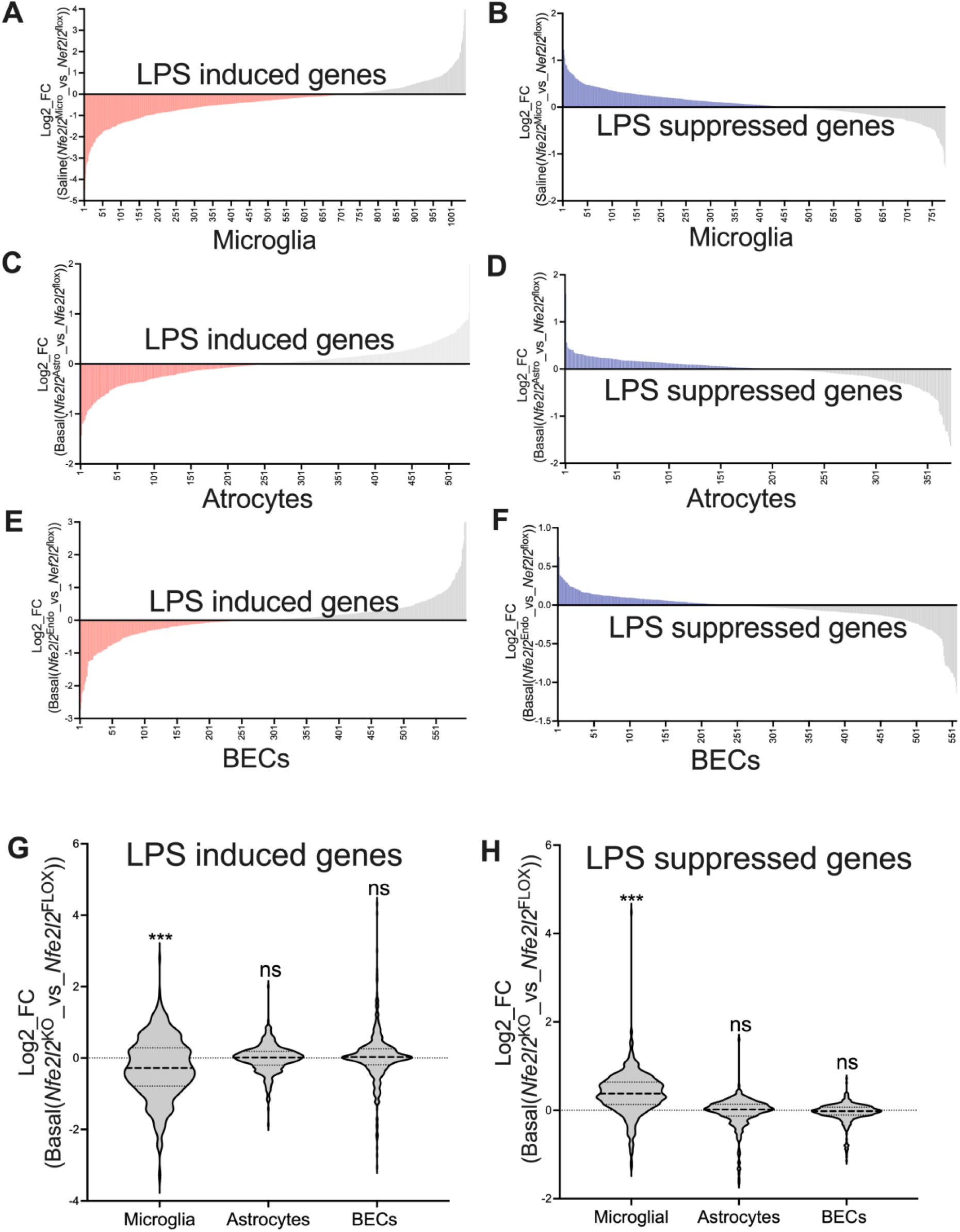
Nrf2 mediates proinflammatory gene expression in microglia, but not in astrocytes and BECs at basal condition. **(A)** The influence of Nrf2 KO in microglia on the expression of microglial LPS-induced gene set at basal condition. For microglial LPS-induced genes (x-axis, FPKM>1, *DESeqP*_adj<0.05, Log_2_FC >1) in WT mice, the Log_2_FC (y-axis) of each gene (x-axis) due to Nrf2 KO in microglia at basal condition were plotted. **(B)** The influence of Nrf2 KO in microglia on the expression of microglial LPS-repressed gene set at basal condition. For microglial LPS-repressed genes (x-axis, FPKM>1, *DESeqP*_adj<0.05, Log_2_FC >1) in WT mice, the Log_2_FC (y-axis) of each gene (x-axis) due to Nrf2 KO in microglia at basal condition were plotted. **(C)** The influence of Nrf2 KO in astrocytes on the expression of astrocyte LPS-induced gene set at basal condition. For astrocyte LPS-induced genes (x-axis, FPKM>1, *DESeqP*_adj<0.05, Log_2_FC >1) in WT mice, the Log_2_FC (y-axis) of each gene (x-axis) due to Nrf2 KO in astrocytes at basal condition were plotted. **(D)** The influence of Nrf2 KO in astrocytes on the expression of astrocyte LPS-repressed gene set at basal condition. For astrocyte LPS-repressed genes (x-axis, FPKM>1, *DESeqP*_adj<0.05, Log_2_FC >1) in WT mice, the Log_2_FC (y-axis) of each gene (x-axis) due to Nrf2 KO in astrocytes at basal condition were plotted. **(E)** The influence of Nrf2 KO in BECs on the expression of BEC LPS-induced gene set at basal condition. For BEC LPS-induced genes (x-axis, FPKM>1, *DESeqP*_adj<0.05, Log_2_FC >1) in WT mice, the Log_2_FC (y-axis) of each gene (x-axis) due to Nrf2 KO in BECs at basal condition were plotted. **(F)** The influence of Nrf2 KO in BECs on the expression of BEC LPS-repressed gene set at basal condition. For BEC LPS-repressed genes (x-axis, FPKM>1, *DESeqP*_adj<0.05, Log_2_FC >1) in WT mice, the Log_2_FC (y-axis) of each gene (x-axis) due to Nrf2 KO in BECs at basal condition were plotted. **(G)** From left to right, *p=5.0E-12, F (1,7700) =47.82 relates to main effect of Nrf2-KO on the expression of microglial LPS-induced gene set under inflammatory condition, 2-way ANOVA; p=0.3441, F (1,4761) =0.8952 relates to main effect of Nrf2-KO on the expression of astrocyte LPS-induced gene set under inflammatory condition, 2-way ANOVA; p=0.4823, F (1,5990) =0.4936 relates to main effect of Nrf2-KO on the expression of BEC LPS-induced gene set under inflammatory condition, 2-way ANOVA **(H)** From left to right, *p=6.0E-09, F (1,3885) =34.00 relates to main effect of Nrf2-KO on the expression of microglial LPS-repressed gene set under inflammatory condition, 2-way ANOVA; p=0.3791, F (1,3357) =4.311 relates to main effect of Nrf2-KO on the expression of astrocyte LPS-repressed gene set under inflammatory condition, 2-way ANOVA; p=0.4693 F (1,6126) =0.5236 relates to main effect of Nrf2-KO on the expression of BEC LPS-repressed gene set under inflammatory condition, 2-way ANOVA

## Discussion

Previous studies have pointed to multiple cytoprotective functions of Nrf2 in the brain^1^, however, cell-type specific role of Nrf2 in adult brain was not clear. Using cell-type specific deletions of Nrf2 in adult mouse brain, our study found that Nrf2 controls expression of slightly overlapping, but mainly distinct, set of genes in microglia, astrocytes and BECs, suggesting that Nrf2 controls distinct transcriptional programmes in different brain cell types. This may be at first glance surprising, particularly for genes positively regulated by Nrf2, since these genes often contain ARE elements that in theory should be responsive to Nrf2 in all cell types. However, not all ARE-containing genes are Nrf2 responsive in all cell types since Nrf2 deletion has different effects on gene expression in different cell types. Some Nrf2 target genes may be epigenetically silenced in certain cell types, making their promoters inaccessible to Nrf2. Moreover, since Nrf2 target genes can be controlled by other transcription factors (such as AP-1)^17,18^ and so the impact of Nrf2 deficiency may depend on the expression of other transcription factors in that cell type.

Nrf2 Global Nrf2 KO mice are susceptible to various pathogen infections^19,20,21,22^ suggesting a protective role of Nrf2 in inflammatory and infectious diseases. However, other studies have suggested a pathogenic role of Nrf2 in certain disorders, including metabolic disorders associated with chronic inflammation, with certain aspects of Nrf2 function (e.g. in lipid metabolism) outweighing the effects of other functions (e.g. antioxidant production)^23,24,25^. Our study showed that Nrf2-deficient microglia had reduced expression of genes involved in responses to oxidative stress and cytokine stimulation, and Nrf2-deficient astrocytes had reduced expression of genes involved in classical pathways of inflammation including MAPK signalling pathway, JAK-STAT signalling pathway, and p53 signalling pathway. In line with this functional categorisation, comparisons of our RNAseq data show that Nrf2-deficiency in microglia and in astrocytes caused a decrease in overall inflammatory responses to a peripheral LPS insult. Taken together, our study suggests that endogenous Nrf2 activity partly mediate responses in microglia and astrocytes to inflammatory conditions, but not in BECs.

It is therefore possible that Nrf2 plays a pro-inflammatory role under certain conditions, although the precise mechanism behind this warrants further investigation. Indeed, a certain level of Nrf2 activity is beneficial and essential in supporting inflammatory responses and controlling pathogenic processes in inflammatory and infectious diseases. Moreover, the absence of Nrf2 may lead to an insufficient inflammatory response, leading to susceptibility to pathogens in global Nrf2 KO mice, whereas in chronic inflammatory diseases such as metabolic disorders, a sustained high level of Nrf2 becomes pathological. Therefore, it is possible that Nrf2’s role in inflammatory responses, and the consequences of these responses, is highly sensitive to both the cell type and the nature of the inflammatory insult.

The fact that Nrf2 activity contributes to the microglial and astrocytic responses to an inflammatory insult is at first glance paradoxical given that enhancing it has been proposed to be a therapeutic strategy^,7,8,9^. Indeed, our recent study also found that pharmacological activation of Nrf2 using a potent Nrf2 activator (RTA-404) suppressed inflammatory responses in microglia, astrocytes and BECs, as well as suppressing leukocyte brain infiltration in response to peripheral LPS insults^11^. Several other studies using Nrf2 activators of natural or synthetic compounds have also supported the notion that pharmacological activation of Nrf2 can limit inflammation in pre-clinical models. In addition, Nrf2 activators, Tecfidera (dimethyl fumarate) has been approved for treating multiple sclerosis^26,27^ and Omaveloxolone (RTA-408) has been approved for treating Friedreich’s Ataxia^28,29^, both potentially in part based on an anti-inflammatory role of Nrf2. Therefore, it is possible that endogenous Nrf2 activity and pharmacological activation of Nrf2 work via different mechanisms and pathways, potentially due to the degree to which they are activated.

To conclude, further studies are required to fully understand the mechanism by which Nrf2 supports inflammatory responses in a cell type-specific manner, enabling any therapeutic strategy to target the right cell at the right time to combat disorders of neuroinflammation.

## Methods

### Mice

The animals involved in this study were all on C57BL/6 background. The Nrf2^*flox*^ mouse line (Mouse Strain Number – 025433), CX3CR1^*CreERT2*^ mouse line (Mouse Strain Number – 031008) and Aldh1l1^*CreERT2*^ mouse line (Mouse Strain Number – 025433) were imported from Jackson Laboratories. The Nrf2^flox^ mice carrying *loxP* sites flanking exon 5 of the *Nfe2l2* gene were crossed to the CX3CR1^*CreERT2*^ mouse and Aldh1l1^*CreERT2*^ mouse to generate *Cx3cr1*^*CreERT2*^: *Nefe2l2*^*fl/fl*^ mice and *Aldh1l1*^*CreERT2*^: *Nefe2l2*^*fl/fl*^ mice respectively. Male mice have been used throughout the study. All procedures described were performed at the University of Edinburgh in compliance with the UK Animals (Scientific Procedures) Act 1986 and University of Edinburgh regulations and carried out under project license numbers P2262369. Mice were group-housed in environmentally enriched cages within humidity and temperature-controlled rooms, with a 12-h light dark cycle with free access to food and water. Mouse genotypes were determined using real-time PCR with transgene specific probes (Transnetyx, Cordova, TN) unless otherwise stated.

### Tamoxifen and LPS Treatments

For induction of Cre recombinase activity, 6-8-week-old *Cdh5*^*CreERT2*^: *Nefe2l2*^*fl/fl*^ mice were given orally with 8 mg tamoxifen (TAM, T5648, Sigma-Aldrich, UK) solved in corn oil (C8267, Sigma-Aldrich, UK) at 4 time points with 24hrs apart. For all experiments, littermates carrying the respective loxP-flanked alles but lacking expression of Cre recombinase (+/+ TAM) were used as controls. Male mice have been used throughout the study. The *Cx3cr1*^*CreERT2*^: *Nefe2l2*^*fl/fl*^ mice, *Aldh1l1*^*CreERT2*^: *Nefe2l2*^*fl/fl*^ mice, and the respective control mice were injected peritoneally with 3mg/kg of LPS (tlrl-3pelps, Invivogen, US) for 24 hrs.

### Isolation of brain cells, Cell Sorting, and RNA extraction

Single brain cells were isolated using Adult Brain Dissociation kit (Miltenyie, 130-107-677, Germany) as per the manufacturer’s instructions. The single brain cell suspension was then incubated with antibodies against CD31 (Biolegend, 102523, UK) CD11b (Biolegend, 101205, UK), ACSA2 (Miltenyie, 130-116-245, Germany), O4 (Miltenyie, 130-117-357, Germany), CD45 (Biolegend, 103125, Germany) for 15 minutes in ice, washed with PBS and then immediately FACS sorted into RNAprotect Cell Reagent (Qiagen, 76526, UK) under the gates of CD31^+^CD45^-^ ACSA2^-^ for BECs, LY6G^-^CD11b^+^CD45^low^ for microglia, ACSA^+^O4^-^ for astrocytes and O4^+^ for oligodendrocytes (Fig. S1). RNA extraction was carried out using the RNeasy Plus Micro Kit (Qiagen) as per the manufacturer’s instructions.

### Quantitative RT-PCR

cDNA was generated using the Transcriptor First Strand cDNA Synthesis Kit (Roche). Seven microlitres of RNA was added to the RT and buffer mixture prepared with random hexamers and oligoDT primers as per kit instructions, and qRT-PCR carried out with the following programme: 10 min at 25 °C, 30 min at 55 °C and 5 min at 85 °C. qPCRs were run on a Stratagene Mx3000P QPCR System (Agilent Technologies) using SYBR Green MasterRox (Roche) with 6 ng of cDNA per well of a 96-well plate, using the following programme: 10 min at 95 °C, 40 cycles of 30 s at 95 °C, 40 s at 60 °C and 30 s at 72 °C, with a subsequent cycle of 1 min at 95 °C and 30 s at 55 °C ramping up to 95 °C over 30 s (to measure the dissociation curve).

### RNA-seq and Analysis

For RNA-seq analysis, libraries were prepared by Edinburgh Genomics using the Illumina TruSeq stranded mRNA-seq kit, according to the manufacturer’s protocol (Illumina). The libraries were pooled and sequenced to 75 base paired-end on an Illumina NovaSeqTM 6000 to a depth of approximately 50 million paired-end reads per sample. For read mapping and feature counting, genome sequences and gene annotations were downloaded from Ensembl version 94. Differential expression (DGE) analysis on data sets was performed using DESeq2 (R package version 1.18.1) using a significance threshold set at a Benjamini-Hochberg-adjusted p-value of 0.05.

## Resource availability

### Lead Contact

Further information and requests for resources and reagents should be directed to and will be fulfilled by the Lead Contact, Jing Qiu.

### Materials Availability

This study did not generate new unique reagents.

### Data and Code Availability

All RNA-seq data that support the findings of this study is available at the European Bioinformatics Institute (ArrayExpress: E-MTAB-14572). All other data are available from the lead contact upon reasonable request.

## Acknowledgements

We would like to thank the Flow Cytometry Facility, led by Dr. Fiona Rossi, at the Centre for Regenerative Medicine at the University of Edinburgh for their help with the Flow Cytometry work. The research leading to these results received funding from the Ann Rowling Regenerative Neurology Clinic at the University of Edinburgh and UK DRI Edinburgh Centre.

## Author contributions

J.Q. conceived the study, performed the experiments, analyzed the data and wrote the manuscript. O.D. and X.H. analysed the data.

